# Jag2 patterns early differentiation in the epidermal stem cell layer

**DOI:** 10.64898/2026.02.28.708758

**Authors:** Sophie Viala, Vandana Nathan, Jacinthe Sirois, Olivia Costanzo, Daniela Perez Laguna, Mauna Musulchi, Milan Heck, Anissa Mourali, Michael H. Mouradian, Katie Cockburn

## Abstract

Adult stem cell function depends on continuous input from the surrounding microenvironment. However, the cues that pattern this function in high turnover tissue that undergo continuous cellular remodeling remain poorly understood. Here we demonstrate that inputs from epithelial neighbors at distinct stages of differentiation cooperatively influence cell fate in the skin epidermis. Stem cells initiate differentiation of their neighbors via the Notch ligand Jag2, linking fate decisions to the composition of the local stem cell environment. When this signal is lost, misoriented divisions and engagement with distinct ligands in the suprabasal layers can partially restore upward flux differentiating cells. Although these mechanisms help to sustain barrier function, they ultimately disrupt tissue architecture, underscoring the need for balanced fates in the stem cell compartment. Together, our findings demonstrate that distinct cellular environments reinforce early epidermal differentiation, with stem cells themselves acting as key mediators of cell fate and tissue organization.

## INTRODUCTION

Stratified epithelia share a common polarized organization, with terminally differentiated cells shed at the tissue surface and replaced by stem cells located several layers away in more internalized cell niches^1–5^. This physical separation necessitates precise coordination so that differentiation events are initiated with the appropriate frequency in the stem cell compartment and proceed with the correct timing as cells transit from one layer to the next. How epithelial cells at distinct stages of differentiation communicate to maintain tissue function in the face of cellular turnover remains a central question in regenerative biology.

In the mammalian skin epidermis, stem cells reside in an underlying basal layer that serves as a site for both self-renewing divisions as well as the initiation of epidermal differentiation^6–8^. During a single day of homeostatic turnover, approximately 5-10% of keratinocytes in the basal layer initiate differentiation^9,10^, a process that involves coincident upregulation of suprabasal structural and adhesive components, downregulation of integrin expression, and gradual upward delamination into the spinous layer directly above^9,11,12^. Increased demand for differentiating cells when the epidermal barrier is compromised can lead to a corresponding increase in the number of basal cells that initiate differentiation^9,10,13,14^, suggesting that this process is under tight environmental control. However, the signals that cue individual cells in the basal layer to initiate differentiation at specific times and places are not well understood.

The contact-based Notch signaling pathway has long been appreciated to play a central role in epidermal differentiation^15,16^. During embryonic development, blocking pathway activity via deletion of Notch 1 and 2 or their obligate downstream transcriptional partner Rbpj prevents early differentiation, leading to a markedly thinner epidermis that lacks an obvious spinous compartment^17,18^. Although the role of Notch signaling in later epidermal maintenance has been less closely studied, early differentiating basal cells as well as spinous cells in adult skin express the canonical Notch target *Hes1*^9,19–21^. Together with the observation that *NOTCH* mutations accumulate in aging human epithelia in part by altering rates of differentiation^22–26^, these data support a continued role for Notch signaling in the patterning of basal cell fate decisions. Early differentiating keratinocytes share a local environment with a range of cell types including fibroblasts, immune cells and keratinocytes both within and across epidermal layers. Each of these populations have the potential to shape Notch activity through the presentation of distinct combinations of ligands^11,27^. However, the specific cell type(s) and relevant ligands that pattern Notch signaling as individual epidermal cells and their neighbors dynamically proliferate, differentiate, and migrate suprabasally have not yet been identified.

Here, we combine genetic loss-of-function models with intravital imaging to interrogate how the local epidermal environment coordinates early stages of keratinocyte differentiation in the adult skin. We show that this process is initiated and maintained by distinct Notch ligands as cells change position, with stem cell-derived Jagged2 (Jag2) driving initial stages of differentiation in the basal layer. In the absence of this key early signal, Notch signaling and differentiation can be restored in cells that reach the suprabasal layers through misoriented cell divisions, but this compensatory behavior leads to defects in tissue architecture over time. Together, our work demonstrates that distinct cellular environments shape and reinforce early differentiation in the epidermis, with stem cells themselves acting as key mediators of cell fate and tissue organization.

## RESULTS & DISCUSSION

### Jag2 activates Notch signaling in the basal layer

To understand how Notch signaling is activated in the adult epidermis, we first examined the expression patterns of known pathway ligands. Transcriptomic data indicate that three ligands, *Dll1*, *Jag1*, and *Jag2*, are produced in distinct but overlapping keratinocyte subpopulations that correspond to different stages of differentiation^5,11,21,28^. To visualize these patterns in intact tissue we performed fluorescence *in situ* hybridization on sagittal sections of adult ear skin and observed that while *Jag1* is expressed broadly throughout both the basal and suprabasal (spinous and granular) layers, *Jag2* and *Dll1* transcripts are mainly present in the basal layer (Figure 1A and 1B). We also noted the occasional presence of underlying stromal cells expressing one or more ligands (Figure 1B, asterisk), indicating that basal cells encounter multiple combinations of Notch ligands from cellular sources residing above, beside and below them.

**Figure 1.**
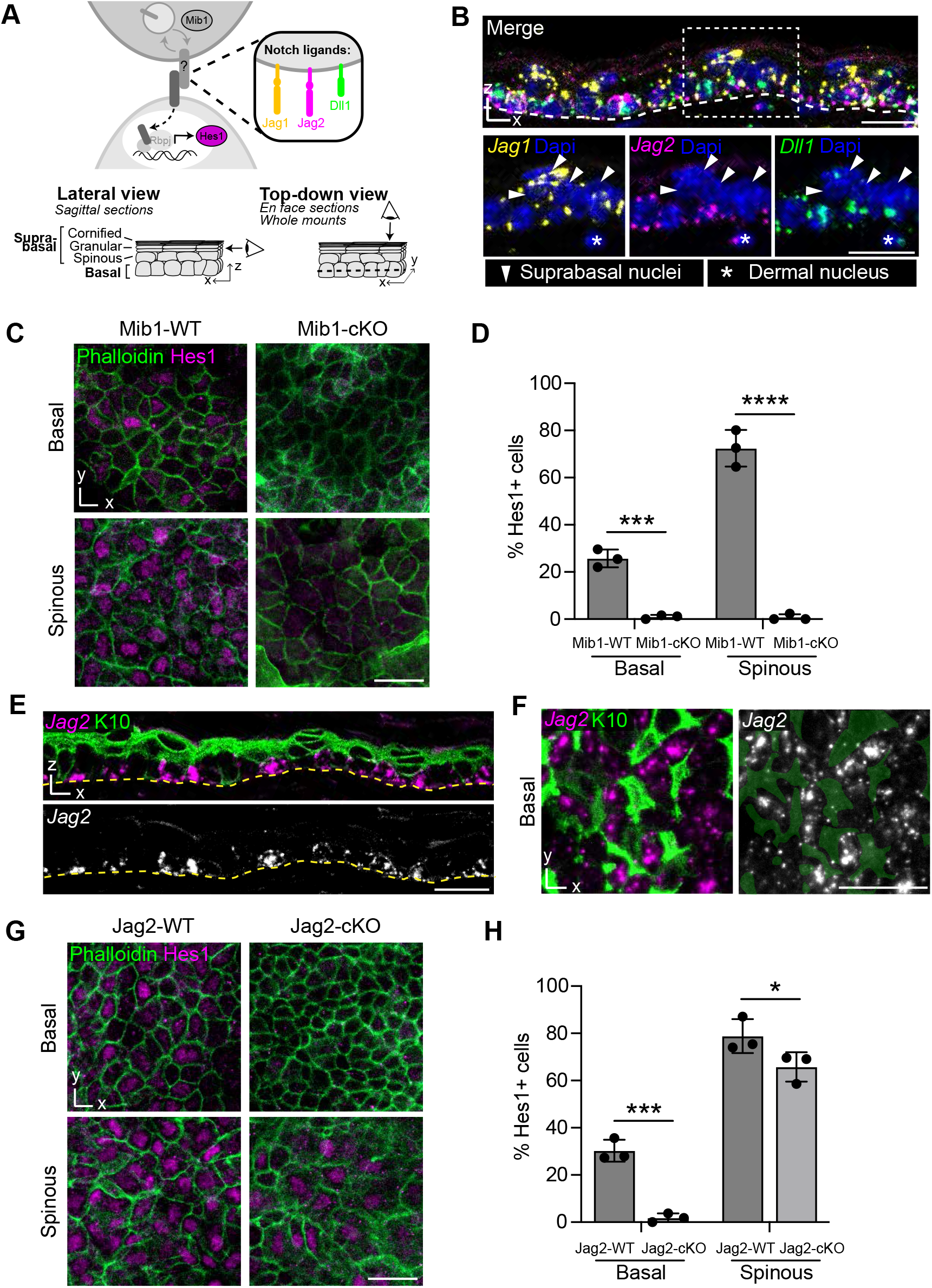
Jag2 activates Notch signaling in the basal layer. (A) Schematic of key Notch ligands and approaches to visualize them in epidermal tissue. (B) Fluorescent in situ hybridization (RNAscope) of Notch ligands Jag1 (magenta), Jag2 (yellow) and Dll1 (green) with Dapi (blue) in a sagittal section of wildtype ear epidermis. Thick white dashed line represents the basement membrane, arrowheads denote suprabasal nuclei, and asterisk indicates a dermal nucleus. (C) Whole-mount staining of Hes1 (magenta) in the basal and spinous layers of Mib1-WT and Mib1-cKO ear epidermis. Cell boundaries are visualized with phalloidin (green). (D) Percentage of basal or spinous cells that express Hes1 in Mib1-WT and Mib1-cKO tissue. Student’s t-test, p≤0.001 (WT vs cKO basal and WT vs cKO spinous). (E) Dual fluorescent in situ hybridization of Jag2 mRNA (magenta or white) and immunostaining of K10 protein (green) in a sagittal section of wildtype ear epidermis. Yellow dashes represent the basement membrane. (F) Dual fluorescent in situ hybridization of Jag2 mRNA (magenta or white) and immunostaining of K10 protein (green) in the basal layer of an en face section of wildtype ear epidermis. Green pseudocolor in right image represents outlines of K10 positive basal cells. (G) Whole-mount staining of Hes1 (magenta) in the basal and spinous layers of Jag2-WT and Jag2-cKO ear epidermis. Cell boundaries are visualized with phalloidin (green). (H) Percentage of basal or spinous cells that express Hes1 in Jag2-WT and Jag2-cKO tissue. Student’s t-test, p≤0.001 (WT vs cKO basal) and p≤0.05 (WT vs cKO spinous). For images in B, C, E, F and G, Scale bars = 20µm. Bar graphs in D and H represent averages from n=3 mice per genotype, 3 regions per mouse, error bars are mean +/- SD.

To first broadly evaluate the role of keratinocyte-derived ligands in epidermal Notch signaling, we performed conditional deletion of *Mindbomb1* (*Mib1*), an E3 ubiquitin ligase essential for the normal endocytic processing of all Notch ligands^29^ (*K5CreERT2; Mib1^flox/flox^*, or Mib1-cKO mice). Wholemount staining and top-down visualization of wildtype adult tissue revealed that the Notch effector gene Hes1 is expressed within a subset of basal cells in a salt-and-pepper fashion that overlaps with the early differentiation marker Keratin 10 (K10), becomes more broadly expressed throughout the spinous population, and is no longer visible once cells reach the granular layers, consistent with previous observations from embryonic skin^18^ (Figures S1A, S1B, 1C and 1D,). Although Mib1-cKO mice died rapidly after Tamoxifen administration, likely due to K5CreER activity in other tissues, we observed that even 5 days of deletion was sufficient to entirely eliminate Hes1 signal from both the basal and suprabasal populations (Figures 1C, 1D and S1B). Thus, epidermal keratinocytes act as the main signal-sending cells that drive Notch activation in the basal and spinous layers.

We next sought to determine which specific ligand(s) are essential for the induction of Hes1 in the basal layer. *In situ* hybridization using epidermal tissue sectioned either sagittally (Figure 1E) or e*n face* to provide a top-down view of the basal layer (Figure 1F) revealed that *Jag2* transcripts are largely absent from K10 positive basal cells. This unique expression pattern, together with evidence that exposure to Jag2 in *trans* can drive keratinocyte differentiation *in vitro*^30^, suggested a possible sender-receiver relationship between basal neighbors at different stages of differentiation. Strikingly, and in contrast to the complete loss of epidermal Notch signaling we observed in Mib-cKO mutants, conditional deletion of Jag2 (*K14CreERT2; Jag2^flox/flox^*, or Jag2-cKO mice; Figure S1C) eliminated Hes1 signal in basal cells but caused only a mild decrease in the number of Hes1 positive cells in the suprabasal layer above, even several weeks after Tamoxifen administration (30 days post-Jag2 deletion; Figures 1G, 1H and S1B). As in control tissue, *Jag1* remained the predominant ligand expressed in the suprabasal layers of Jag2-cKO tissue (Figure S1C), suggesting a potential role for this ligand in the maintenance of Notch activity once cells exit the basal compartment. These results indicate that Notch signaling is patterned by distinct ligands as cells move upward through the epidermal layers, with stem cell-derived Jag2 triggering initial pathway activity in the basal layer and other keratinocyte-derived ligands such as Jag1 maintaining activity in the spinous layer.

### Loss of Notch activity in the basal layer delays epidermal differentiation

We next assessed how an absence of Notch signaling in the basal layer altered epidermal differentiation in Jag2-cKO mutants. Consistent with the loss of Hes1 signal in this compartment, wholemount staining revealed that Jag2-cKO tissue after 30 days of deletion contained only a small, scattered population of K10-positive basal cells (Figures 2A and 2B). To assess how this decrease in the early differentiating population affected maintenance of the suprabasal layers, we examined tissue organization in sagittal sections. Collectively, the suprabasal compartment was thinner in mutant epidermis (Figures 2C, 2D, and S2A), due to a reduction in both the K10 single-positive spinous population as well as the overlying K10/involucrin double-positive granular layers (Figure S2B). To test whether this thinner granular compartment formed tight junctions and maintained a functional inside-out barrier, we assessed the diffusion of an intradermally injected NHS-LC-biotin tracer^31^. As in control epidermis, intercellular biotin diffused upward after injection but was halted at regions of apical Claudin-4 accumulation (Figure S2C), indicating the presence of a functional tight junction network between granular cells in Jag2-cKO mutants. Thus, despite a decrease in the number of early differentiating cells produced in the basal layer of Jag2-cKO mutants, the tissue remains able to generate a sufficient number of suprabasal cells to maintain barrier function.

**Figure 2.**
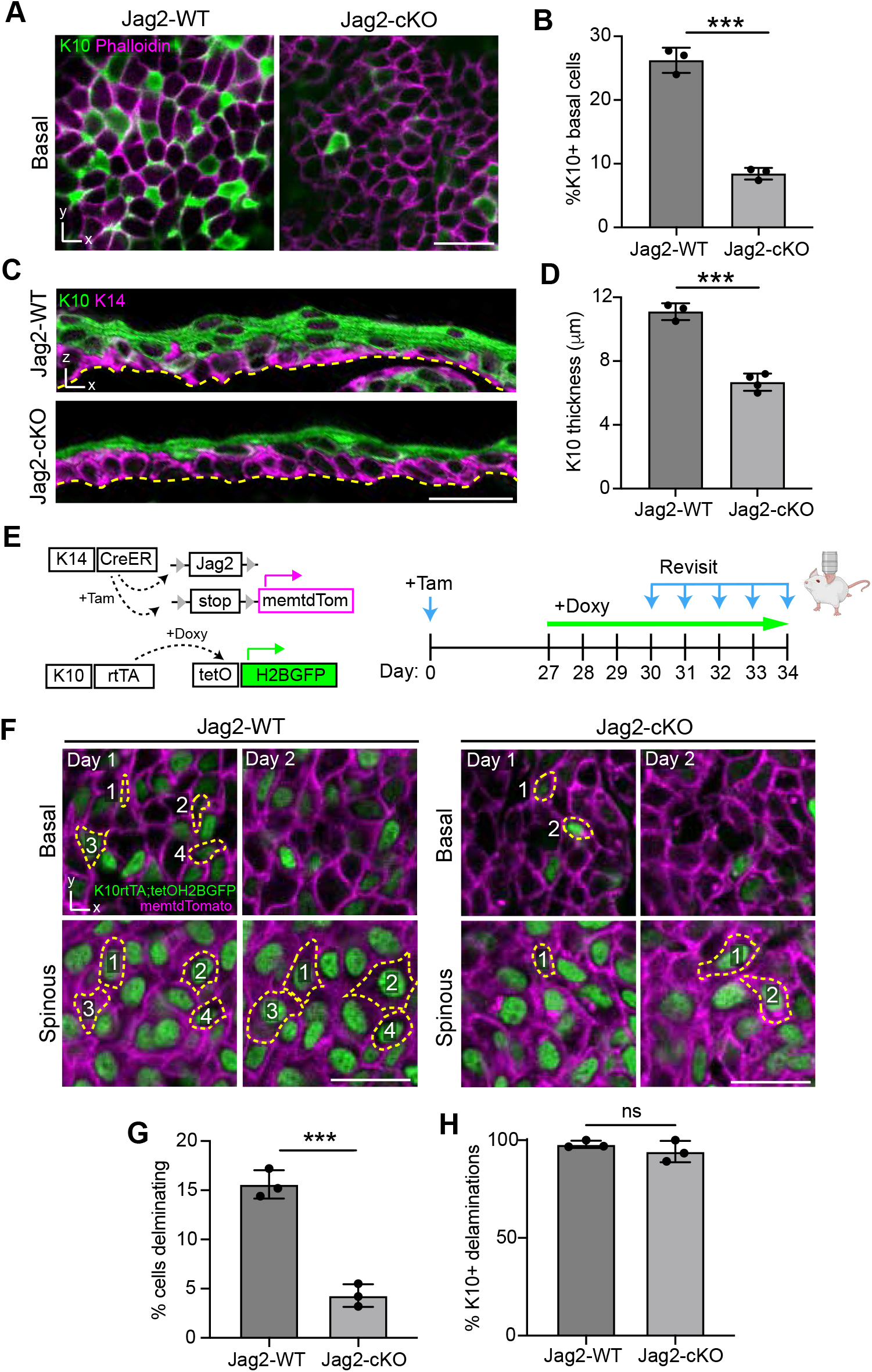
Loss of Notch activity⊠in the basal layer⊠delays epidermal differentiation. (A) Whole-mount staining of K10 (green) in the basal layer of Jag2-WT and Jag2-cKO ear epidermis at 30 days post-deletion. Cell boundaries are visualized with phalloidin (magenta). (B) Percentage of basal cells that express K10 in Jag2-WT and Jag2-cKO tissue. Student’s t-test, p≤0.001 (C) Immunostaining of K10 (green) and K14 (magenta) in sagittal sections of Jag2-WT and Jag2-cKO ear epidermis at 30 days post-deletion. Yellow dashes represent the basement membrane. (D) Total thickness of Keratin10+ layers in sagittal sections of Jag2-WT and Jag2-cKO ear epidermis. Student’s t-test, p≤0.001. (E) Schematic of experimental design to visualize basal cell delamination. Epidermal regions were longitudinally revisited starting 30 days after Jag2 deletion and 3 days after Doxycycline administration (K10 reporter induction). (F) Representative images of Jag2-WT and Jag2-cKO epidermal regions expressing the K10 reporter (green) revisited on Days 1 and 2 of imaging. Epidermal plasma membranes are marked with tdTomato (magenta). Yellow dotted lines indicate basal cells, often visible across the basal and spinous layers at Day 1, that have lost basement membrane contact and completed delamination on Day 2. (G) Percentage of basal cells completing delamination over 24h in Jag2-WT and Jag2-cKO tissue. Student’s t-test, p≤0.001. (H) Percentage of delaminating basal cells that express the K10 reporter in Jag2-WT and Jag2-cKO tissue. Student’s t-test, p>0.05. For images in A, C, and F, Scale bars = 20µm. Bar graphs in B, D, G and H represent averages from at least n=3 mice per genotype, 3 regions per mouse, error bars are mean +/- SD.

In adult epidermis, early differentiation and delamination out of the basal layer are temporally and molecularly linked, with individual basal cells gradually inducing differentiation-associated transcripts as they transition upward into the spinous layer over the course of several days^9,32^. To determine whether the reduced basal cell differentiation we observed in Jag2-cKO tissue is accompanied by a corresponding reduction in the number of delamination events, we combined our conditional deletion model with a previously established K10 reporter system (*K10rtTA; tetO-H2BGFP*^9,33^) and an inducible fluorescent membrane marker (*iSuRe-HadCre*^34^) (Figure 2E). In resulting mice, epidermal cells can be visualized with membrane-localized tdTomato, and K10-positive cells can be further identified via expression of H2BGFP. Initial assessment of the resulting tissue revealed a marked decrease in the number of K10 reporter-positive basal cells in Jag2-cKO mutants, consistent with the reduction of K10-positive cells we observed via immunostaining (Figures S2D and S2E). We therefore combined this system with longitudinal intravital imaging^9^ to assess cell movement and differentiation status in the basal layer of Jag2-cKO tissue over time (Figure 2E and S2F).

To evaluate the frequency and timing of basal cell delamination in Jag2-cKO epidermis, we performed daily revisits of the same epidermal regions and focused specifically on cells that could be seen losing contact with the basement membrane over a 24h period (Figures 2F and S2F). In agreement with the significant reduction in K10-positive basal cells we observed in both fixed and live tissue, these delamination events were rare in Jag2-cKO mutants (Figures 2F and 2G). The few residual delaminating cells we could observe in mutant tissue were almost exclusively K10 reporter positive, just as we observed in controls (Figure 2H). Notably, revisits as well as timelapse imaging of mutant tissue over multiple hours did not reveal any obvious examples of more rapid cell extrusion events reminiscent of those observed in the developing epidermis and other epithelial systems^35,36^. Our observation that individual cells cannot delaminate or otherwise lose basement membrane contact unless they have initiated differentiation suggests that alternative mechanisms may compensate to maintain the epidermal barrier when differentiation is reduced.

### Perpendicular divisions generate suprabasal cells when basal delamination is reduced

Despite markedly reduced rates of delamination in Jag2-cKO tissue, we found that the basal layer remained highly proliferative and without any significant change in rates of cell death (Figures S3A-S3D). Due to this imbalance in cell loss vs cell gain over time, the basal layer of mutant tissue was significantly denser, with cells appearing tightly packed and more columnar in shape (lateral view in Figure 2C; quantified in Figure S3E). Given the established link between cell density, cell geometry and division orientation in the basal layer^37,38^, we hypothesized that misoriented cell divisions could serve as an alternative mechanism to deposit cells suprabasally in Jag2-cKO tissue. Using both control and Jag2-cKO tissue combined with the fluorescent alleles described above, we performed intravital timelapse imaging and assessed cell division orientation with respect to the basement membrane (Figures 3A-C). In control tissue, as expected^39,40^, the majority of cell divisions occurred parallel to the basement membrane (< 30° orientation), with a smaller subset appearing more oblique (between 30° and 60°) and very rare occurrences of truly perpendicular divisions (>60°) (Figure 3C). In contrast, close to one quarter of cell divisions in Jag2-cKO tissue were fully perpendicular, with several occurring at almost exactly right angles to the basement membrane (Figures 3C). These perpendicular divisions, which occurred regardless of K10 reporter status (Figure S3F), produced daughters that had completely lost contact with the basement membrane and were entirely suprabasal at the end of telophase (Figure 3B).

**Figure 3.**
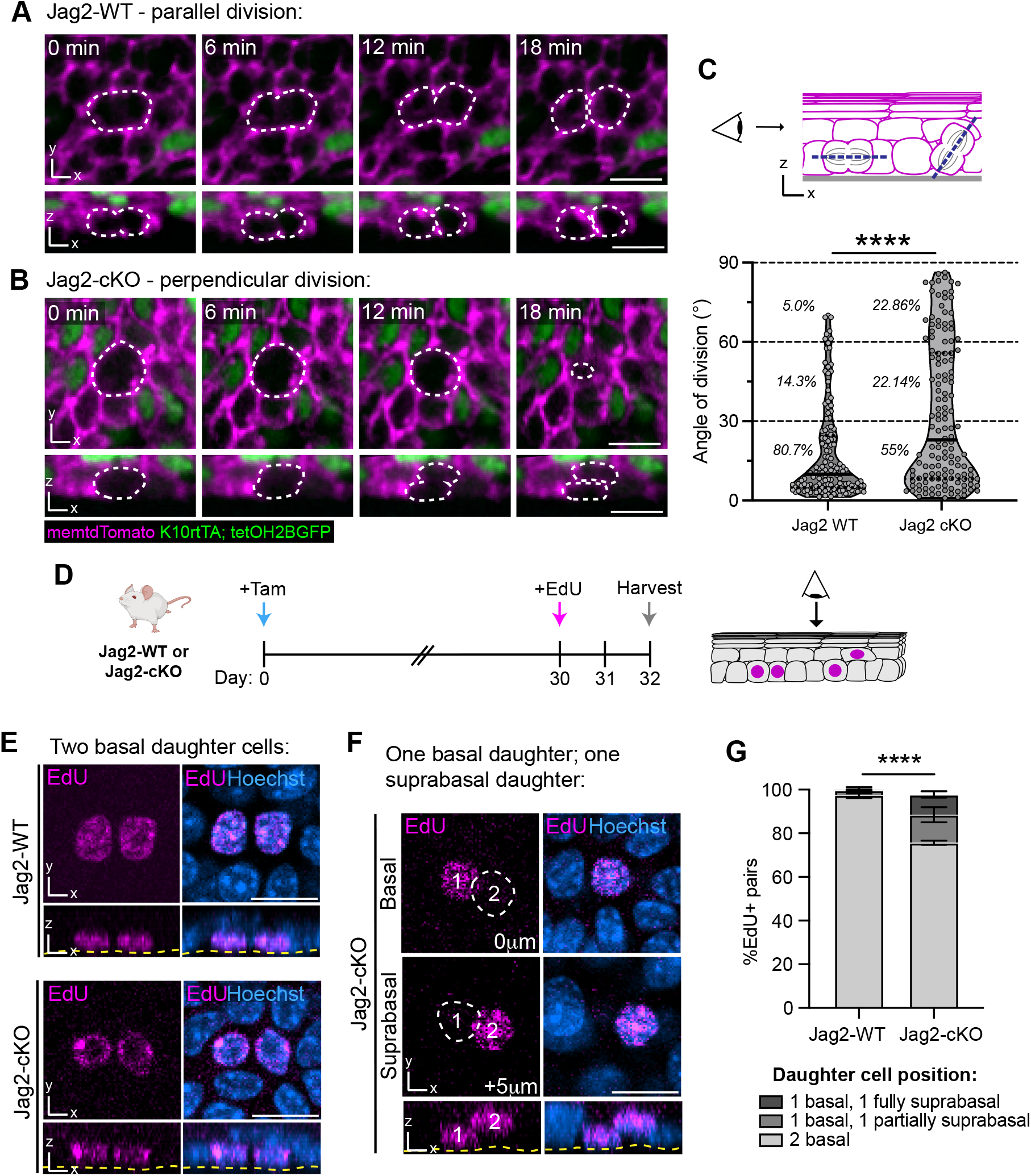
Perpendicular divisions generate suprabasal cells when basal delamination is reduced. (A) Stills from timelapse imaging of Jag2-WT epidermis showing a mitotic cell (dotted white outline) dividing parallel to the basement membrane and yielding two basal daughter cells. (B) Stills from timelapse imaging of Jag2-cKO epidermis showing a mitotic cell (dotted white outline) dividing perpendicular to the basement membrane and yielding one suprabasal daughter cell and one basal daughter cell. Top-down view shown here (top panel) taken at the cleavage furrow region between upper and lower daughters. (C) Quantification of the angle of division of mitotic cells in Jag2-WT and Jag2-cKO tissue at 30 days post-deletion. Proportion of divisions within 0-30°, 30-60° and 60-90° bins indicated in italics. Solid line represents the median; dotted lines represent the quartiles. Kolmogorov-Smirnov test, p<0.0001. Graph represents n=3 mice and 140 divisions per genotype. (D) Schematic of 48h EdU pulse-chase experiment to follow daughter cell localization and fate in Jag2-cKO tissue at 30 days post-deletion. (E) Representative images of EdU+ (magenta) basal daughter pairs in Jag2-WT and Jag2-cKO tissue. Hoechst (blue) marks all nuclei. (F) Representative images of EdU+ (magenta) daughter cells, one basal and one suprabasal, in Jag2-cKO tissue. Hoechst (blue) marks the nuclei. (G) Percentage of EdU+ daughter cell pairs that are symmetrically (both basal) or asymmetrically localized (one basal and one partially/fully suprabasal) at 48h post-EdU incorporation in Jag2-WT and Jag2-cKO tissue 30 days post-deletion. Contingency tests performed per biological replicate and combined using Fisher’s method; p<0.0001. Graph represents n=3 mice, 3 regions per mouse. Error bars are mean +/- SD. For images in A, B, E and F, Scale bars = 20µm.

Because corrective mechanisms both during and after mitosis can influence the final positioning of daughter cells^41–43^, we further assessed the relationship between division angle and later daughter cell position in Jag2-cKO tissue. Although suprabasal daughters generated during timelapses of Jag2-cKO tissue were often difficult to conclusively identify when tissue was revisited at later timepoints, we identified several examples in which progeny retained their spinous location up to 24h later (Figure S3G). To track these events more quantitatively, we administered a single dose of EdU to control and Jag2-cKO mice and harvested tissue 48h later (Figure 3D). In control tissue, newly generated daughter cells remained exclusively basal over this 48h period (Figures 3E and 3G). In contrast, roughly one quarter of Jag2-cKO daughter cell pairs were positionally asymmetric, with one daughter cell nucleus in the basal layer and the other positioned several microns above (Figures 3E, 3F and 3G). We categorized these non-basal daughters as either partially suprabasal (nucleus located between the basal and spinous layers) or fully suprabasal (nucleus fully integrated into the spinous layer) (Figure 3G). The proportion of Jag2-cKO daughter cell pairs that contained a partially or fully suprabasal daughter cell closely matched the proportion of perpendicular divisions we observed via timelapse imaging, indicating that suprabasally deposited daughter cells retain their position well after mitosis is complete. To evaluate the relationship between daughter cell position and fate, we combined EdU labelling with wholemount K10 staining. 48h after EdU incorporation, many partially or fully suprabasal cells in the Jag2-cKO tissue were indeed K10 positive (Figure S3G, H and I), indicating a capacity to differentiate once these cells have moved out of the basal compartment.

### Loss of Notch signaling leads to compromised basal layer organization

To assess whether perpendicular divisions can serve as a long-term mechanism to maintain tissue architecture in the absence of basal Notch signaling, we examined the organization of Jag2-cKO mutants at a later experimental timepoint (55 days post-deletion). Although the number of differentiating basal cells remained low at this timepoint, we observed no additional thinning of the suprabasal layers compared to Jag2-cKO epidermis at 30 days post-deletion (Figures 4A, S4A and S4B). At this later stage however, the stem cell compartment of Jag2-cKO mutants had become hyperplasic, with undifferentiated K14-positive basal cells piling on top of one another to generate a thickened, pseudostratified structure (Figures 4A, 4B and S4C). This hyperplasia became more pronounced over time, with Jag2-cKO tissue at 6 months post-deletion containing a strikingly multilayered K5-positive population underlying a comparatively well-organized K10-positive suprabasal compartment (Figure S4D).

**Figure 4.**
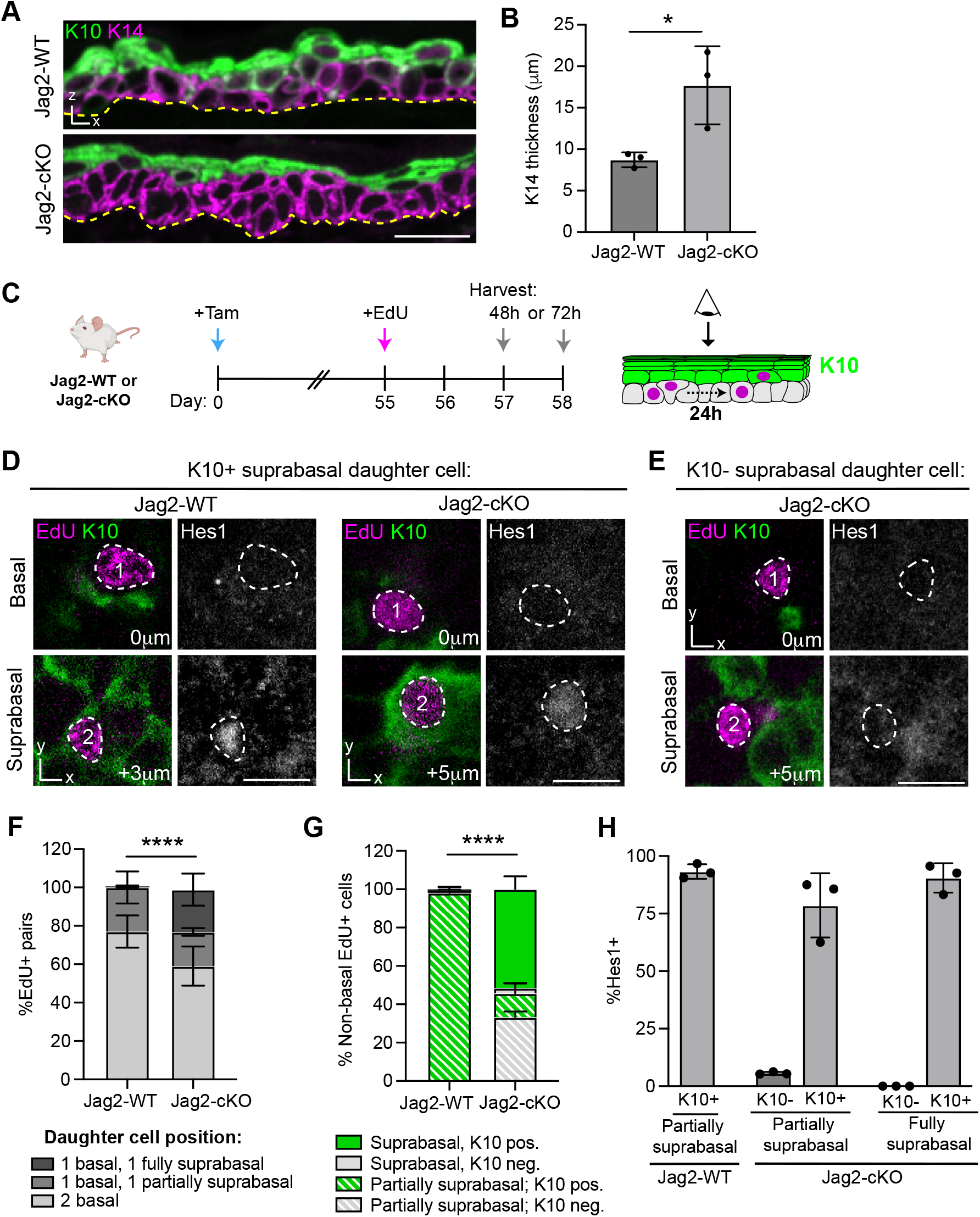
Loss of Notch signaling leads to compromised basal layer organization. (A) Immunostaining of K10 (green) and K14 (magenta) in sagittal sections of Jag2-WT and Jag2-cKO ear epidermis at 55 days post-deletion. Yellow dashes represent the basement membrane. (B) Total thickness of Keratin14+ layers in sagittal sections of Jag2-WT and Jag2-cKO ear epidermis. Student’s t-test, p≤0.05. (C) Schematic of EdU pulse-chase experiment to follow daughter cell localization and fate in Jag2-cKO tissue at 55 days post-deletion. Daughter cell position and K10 expression were compared at 48h vs 72h post EdU administration to assess the differentiation of suprabasal daughters over time. (D) Representative images of asymmetrically localized EdU+ daughter cell pairs (magenta) containing one suprabasal daughter cell that is also K10 and Hes1 positive (green and gray, respectively) in Jag2-WT and Jag2-cKO tissue. (E) Representative image of asymmetrically localized EdU+ daughter cell pair (magenta) containing one basal and one suprabasal daughter cell that are both K10 and Hes1 negative in Jag2-cKO tissue. (F) Percentage of EdU+ daughter cell pairs that are symmetrically (both basal) or asymmetrically localized (one basal and one partially/fully suprabasal) at 72h post-EdU incorporation in Jag2-WT and Jag2-cKO tissue 55 days post-deletion. Contingency tests performed per biological replicate and combined using Fisher’s method; p<0.0001. (G) Percentage of EdU+ daughter cells scored as partially or fully suprabasal in Jag2-WT and Jag2-cKO tissue that express K10. Contingency tests performed per biological replicate and combined using Fisher’s method; p<0.0001. (H) Percentage of Hes1+ cells according to position and K10 expression in Jag2-WT and Jag2-cKO tissue. For images in A, D, E, Scale bars = 20µm. Bar graphs in B, F, G and H represent averages from at least n=3 mice per genotype, 3 regions per mouse, error bars are mean +/- SD.

To better understand this transition towards hyperplasia, we assessed cell division outcomes at 55 days post-deletion by again administering a single dose of EdU and harvesting tissue 48h later (Figure 4C). Consistent with earlier timepoints, roughly one quarter of divisions in Jag2-cKO tissue generated asymmetrically positioned basal/suprabasal daughter pairs (Figure S4E), indicating that perpendicular divisions continue at a similar frequency over time. Many of the daughter cells born at 55 days post-deletion were also capable of initiating K10 expression (Figure S4D). Notably however, a subset of partially suprabasal daughter cells did not express detectable levels of K10 (Figure S4D), suggesting delayed or defective differentiation. To evaluate the former possibility, we repeated our EdU labelling experiment in Jag2-cKO mutants at 55 days post-deletion, this time harvesting tissue 72h after labelling to allow additional time for K10 induction (Figure 4C). This 72h chase was sufficient for us to observe EdU-labelled daughter cells in control tissue that had started to delaminate upwards, and as expected, these partially suprabasal cells had also induced K10 (Figure 4D, 4F and 4G). However despite the extended timeframe, Jag2-cKO tissue retained a population of K10-negative, partially suprabasal daughter cells that appeared unable to differentiate or transition upward (Figure 4E and 4G). We hypothesized that unlike daughter cells deposited directly into the spinous layer and exposed to alternative sources of Notch-activating ligand, these more transitional cells may lack sufficient contact with suprabasal neighbors for pathway activation. Indeed, while partially and fully suprabasal K10-positive daughter cells in Jag2-cKO tissue were Hes1-positive, partially suprabasal K10-negative daughters were almost exclusively Hes1-negative (Figures 4D, 4E and 4H). Together, these results indicate that perpendicular divisions can serve as an alternate mechanism to deposit daughter cells into the suprabasal environment, promoting Notch signaling and differentiation. However, misoriented divisions in which daughter cells are deposited in more intermediate positions may not allow for this cell fate transition to occur.

In this study, we demonstrate that keratinocytes at distinct stages of differentiation shape the fate and position of their neighbors via Notch signaling. Jag2 produced by stem cells is essential to trigger Notch activity, differentiation and delamination of basal neighbors, while distinct ligands such as Jag1 maintain this signal once cells enter the suprabasal layers. Although misoriented divisions can help sustain an upward flux of cells when basal differentiation and delamination are blocked, this eventually compromises tissue organization and architecture of the stem cell compartment. Together, our results highlight the complex feedback that patterns cellular turnover in the epidermis.

Despite longstanding efforts to understand how self-renewal and differentiation are balanced at a tissue-wide level through lineage tracing and transcriptomic analyses of epidermal stem and progenitor cells^10,40,44–47^, the cellular and molecular logic governing the timing and location of individual stem cell differentiation events in the basal layer have remained unclear. Our results indicate that the expression of Jag2 may serve as one such level of control, allowing the relative proportion of undifferentiated (Jag2+) and committed (Jag2-) neighbors to modulate an individual basal cell’s Notch activity level and probability of differentiating. This feedback mechanism could allow cells to dynamically adjust their behaviors as some neighbors delaminate and others divide, initiating differentiation in response to the changing composition of the local basal neighborhood. Recent transcriptomic analyses indicate that *Jag2* is a conserved marker of undifferentiated basal cells across a range of stratified epithelia from diverse germ layer origins^5^, suggesting that the sender-receiver relationship we propose here may represent a more universal mechanism.

In adult epidermis, cell density is maintained in part through tight control of basal cell proliferation. Elimination of basal cells via delamination or death allows neighbors to expand, progress through the cell cycle, and divide, closely balancing cell loss with cell gain^10,20,48^. Our data indicate that this homeostatic coupling is lost when cells in the basal layer are unable to differentiate efficiently, with rates of proliferation remaining steady despite a significant reduction in cells lost through upward delamination. Although the mechanisms behind this uncoupling are not yet clear, the more columnar shape of Jag2 mutant basal cells suggests that they may have an altered capacity to expand vertically as opposed to laterally, potentially due to altered mechanical constraints from the spinous and granular layers above^49^. Expansion along the apical-basal axis may also facilitate the emergence of the misoriented cell divisions that we observed in Jag2 mutant tissue^37^. Although the emergence of perpendicular divisions in stratified adult tissue clearly helps to maintain the cellularity and function of the suprabasal layers, our results also highlight that the preservation of barrier function through altered spindle orientation can occur at the expense of an organized stem cell compartment.

Finally, our work helps to elucidate how a contact-based signaling pathway such as Notch can be maintained despite the continuous upward movement and neighbor exchange taking place during epidermal turnover^9,10,40,50^. Although pathway activity is first triggered by undifferentiated cells in the basal layer, it is maintained by a distinct ligand, likely Jag1, as delaminating cells enter the spinous compartment. This combined input from multiple cell types likely helps to ensure that the transcriptional changes that accompany differentiation are closely matched to the positional changes associated with suprabasal movement^9,12^. Together, our work highlights the extent to which the local keratinocyte environment shapes epidermal differentiation as cells transit from the stem cell compartment toward the skin’s surface.

## ACKNOWLEDGEMENTS

Thank you to members of the Cockburn lab and Drs. Chen Kam, Mélanie Laurin, Yulia Shwartz and Adda-Lee Graham-Paquin for critical feedback on the manuscript. We thank Christopher Glasz and Vikram Nathan for providing Python scripts for tissue thickness measurements. Image acquisition was performed in the McGill University Advanced BioImaging Facility and tissue processing, embedding, and sectioning was performed by the Goodman Cancer Institute Histology Innovation Platform. We thank Plinio Queiroz Da Cruz for help optimizing *en face* sectioning for paraffin-embedded epidermal tissue. The following investigators kindly provided mouse lines used in this study: Jayaraj Rajagopal (*Jag2^flox^* and *Mib1^flox^*), Rui Benedito (*iSuRe-HadCre*), and Terry Lechler (*K10rtTA*). This work is supported by grants from the Canadian Institutes for Health Research (PJT-190288), the Natural Sciences and Engineering Research Council of Canada (RGPIN-2023-04161), and the Richard & Edith Strauss Foundation to K.C. V.N. and D.P.L. received support through Master’s scholarships from the Fonds de recherche du Québec—Santé. M.H. was supported by a Défi-Canderel Entry Scholarship from the Rosalind and Morris Goodman Cancer Institute and holds a Masters’ training scholarship from the Fonds de recherche du Québec—Nature et technologies. O.C. was supported by an Undergraduate Student Research Award from the Natural Sciences and Engineering Research Council of Canada. M.M. was supported by a Défi-Canderel Rising Star Award and a Graduate Studentship from the Rosalind and Morris Goodman Cancer Institute and a McGill Faculty of Medicine and Health Sciences Internal Scholarship. K.C. is a Tier 2 Canada Research Chair in Dynamics of Tissue Regeneration.

## AUTHOR CONTRIBUTIONS

Conceptualization: S.V. and K.C. Methodology: S.V., V.N., J.S., M.H., M.H.M., and K.C. Investigation: S.V., V.N., J.S., O.C., M.M., D.P.L., A.M., and K.C. Formal analysis: S.V., V.N., O.C., M.M., D.P.L., and K.C. Software: S.V. and V.N. Validation: S.V., V.N., O.C., M.M., D.P.L., and K.C. Data curation: S.V., V.N., and K.C. Funding acquisition: K.C. Project administration: S.V., V.N. and K.C. Supervision: K.C. Writing – original draft: S.V. and K.C. Writing – review & editing: all authors.

## MATERIALS AND METHODS

### Mice and experimental treatments

*Jag2^flox^* and *Mib1^flox^* (Koo et al, 2007) mice^29,51^ were obtained from J. Rajagopal (Harvard). *K10rtTA* mice^33^ were obtained from V. Greco (Yale). *iSuRe-HadCre* mice^34^ were obtained from R. Benedito (CNIC). *K14CreERT2*^52^, *K5CreERT2*^53^, and *tetO-H2BGFP*^54^ were obtained from the Jackson Laboratory (strains #038390, #029155 and #005104, respectively). Mice were maintained on a CD1 background (*Jag2^flox^;K14CreERT2*) or CD1/C57Bl/6J mixed background (*Mib1^flox^;K5CreERT2*). To delete *Jag2* or *Mib1* in adult epidermis, 6- to 8-week-old mice received daily intraperitoneal injections of 100µl (for 5 days) or 150µl (for 3 days) of 20mg/mL tamoxifen in corn oil. Controls were *K14CreERT*2 or *K5CreERT2* mice with wildtype *Jag2* and *Mib1* alleles subjected to the same tamoxifen regimen as their cKO counterparts. To visualize *Krt10* promoter activity and epidermal cell membranes, Jag2^flox^; K14CreERT2; iSuReHadCre; K10rtTA; tetO-H2BGFP received daily intraperitoneal injections of 100µl 20mg/mL tamoxifen (for 5 days) followed by doxycycline water (2mg/ml) in drinking water with 2% sucrose continuously starting 3 days before intravital imaging experiments. To assess proliferation rates, mice were injected intraperitoneally with EdU (50µg/g in PBS) and sacrificed 2h later. To assess cell fate post-mitosis, mice were injected intraperitoneally with EdU (25µg/g in PBS) and sacrificed 48h or 72h later. Mice from the experimental and control groups were randomly selected from either sex. All animal procedures were carried out in accordance with the Canadian Council on Animal Care guidelines and approved by the McGill University Animal Care Committee (#MG10013).

### Wholemount staining

To isolate epidermis for whole-mount staining, ear skin was mechanically separated from the cartilage using forceps and floated over 5 mg/mL dispase II solution (reconstituted in PBS) at 37°C for 15 minutes. The epidermis was separated from the dermis using forceps and fixed in 4% paraformaldehyde in PBS for 20 min (Hes1 staining) or 45 minutes (all other stainings) at room temperature, washed with PBS, permeabilized and blocked (0.2% Triton-X, 5% NDS and 1% BSA in PBS) for 2 hours at room temperature or overnight at 4°C. Samples were then incubated in primary antibody mix in blocking solution overnight at 4°C. Following this incubation, samples were washed 3 times for 10 min in blocking solution, incubated with secondary antibodies in blocking solution for 1h30 at room temperature, washed 3 times for 10 min in blocking solution and mounted with anti-fade mounting medium (90% glycerol, 10% PBS, 0,2% n-propyl gallate, 0,1% sodium azide). To visualize EdU incorporation, epidermal tissue was isolated and washed twice in a 3% BSA in PBS solution. The Click-iT® reaction was performed according to the manufacturer’s protocol, for 30 minutes at room temperature, protected from light (250µl of mix per tissue). The tissues were then washed in 3% BSA in PBS, permeabilized, blocked, and whole mount staining was performed as described above.

### Paraffin-embedded section staining

Samples were fixed for 24h in 4% PFA. Paraffin embedding and sectioning (4µm) was performed on Assure Charged Slides (EPIC Scientific) by the Histology Innovation Platform at the Rosalind and Morris Goodman Cancer Institute. For immunofluorescence, slides were warmed to 50°C, and immersed in HistoClear II twice (5 min and 3 min), followed by 3 min in 100% ethanol, 95% ethanol, and 70% ethanol, which was then gradually changed to dH2O. Antigen retrieval was performed by immersing the slides in Tris-EDTA buffer and incubating them in a high-pressure cooker for 7 min, followed by a 20 min incubation at room temperature and pressure, 2 washes in dH2O and 2 washes in PBS for 5 min. Slides were then blocked 1h in 5%NGS and incubated in primary antibody mix overnight at 4°C in a humid chamber. Slides were washed in PBS and PBST and incubated in secondary antibody mix at room temperature for 1h30, washed again in PBS and PBST, and mounted with anti-fade mounting medium (90% glycerol, 10% PBS, 0,2% n-propyl gallate, 0,1% sodium azide).

### *In situ* hybridization

Fluorescent in situ hybridization (ISH) was performed on paraffin-embedded sections using the RNAscope Multiplex Fluorescent Reagent V2 kit (ACD Bio-Techne) following manufacturer’s instructions. Target retrieval was performed with a 15 minute incubation in a steamer and protease III treatment (322381) was applied to tissues. Sections were hybridized with the following probes: Mm-Jag1 (412831), Mm-Jag2 (417511-C3) and Dll1 (425071-C2), followed by TSA Vivid fluorophores (323270; 1:1000). We used the 3-Plex Positive Control Probe which targets the housekeeping genes POLR2A, PPIB and UBC, and the 3-plex Negative Control Probe which targets the bacterial gene dapB (one probe per detection channel). For dual ISH-IHC with Keratin1 0 antibody, after the last RNAScope Blocker step, slides were rinsed in PBST, followed by 1 hour incubation in blocking buffer (PBS, BSA 1%, NGS 10%) at room temperature followed by standard immunofluorescence staining as described above. Slides were finally mounted using Prolong Diamond Antifade (Life Technologies P36970) and DAPI.

### Epidermal Barrier Assay

EZ-LinkTM Sulpho-N-hydroxysuccinimide-long-chain spacer arm double long chain-Biotin (Sulpho-NHS-LC-LC-Biotin) was reconstituted at 10 mg/mL in PBS containing 10mM CaCl_2_ no more than 30 minutes before use. Following anesthesia with 5% isoflurane, 20µL of biotin solution was injected intradermally into dorsal ear skin using a 31G needle (VWR 76290) inserted at a 20°C angle towards the periphery of the ear. Mice were kept under anesthesia for another 30 minutes to allow for dye penetration. Tissue was then harvested, embedded in Clear Frozen Section Compound, flash frozen in a mix of acetone and dry ice and stored at -80°C. Cryosectioning (10µm) was performed by the Histology Innovation Platform and slides were stored at -80°C. To stain, slides were dried at room temperature for 45 min, followed by a 10 min fixation in 4% PFA and a 5 min wash in PBS. Antigen retrieval was performed by incubating the slides 10 min in PBST + NaCitrate (10mM) + 0.2% Triton-X, followed by a 5 min wash in PBST. The slides were blocked for 1h in 5% NGS and incubated in primary antibody mix (Guinea Pig anti-Keratin 10 1:200, Rabbit anti-Claudin 4 1:100 in PBS) overnight at 4°C. The slides were then washed in PBS and PBST and incubated in secondary antibody mix (Cy3 Streptavidin 1:200, goat anti-rabbit Alexa Fluor 647 1:500, goat anti-guinea pig Alexa Fluor 488 1:500, Hoechst 33342 1:500, 1% Triton-X-100, 2% NGS) at room temperature for 1h30. Slides were washed in PBS and PBST and mounted with anti-fade mounting medium (90% glycerol, 10% PBS, 0,2% n-propyl gallate, 0,1% sodium azide).

### Intravital imaging

All imaging was performed in distal regions of the ear skin during prolonged telogen, with hair removed using depilatory cream (Nair) at least one week before the start of each experiment. Mice were anesthetized with vaporized isoflurane delivered via nose cone. Images were acquired with a Zeiss LSM880NLO microscope equipped with a Chameleon Discovery NX multiphoton laser (Coherent) using a 20x water immersion lens (Zeiss, NA:1.0). Z-stacks were acquired in 1µm stacks covering the entire thickness of the epidermis. To follow the same epidermal cells over multiple days, inherent landmarks of the skin together with a micro-tattoo were used to navigate back to the same epidermal regions every 24h. For timelapse imaging, serial optical sections were obtained every 5 minutes for at least 1h.

### Imaging and quantifications

Images of paraffin-embedded tissue were collected in the McGill University Advanced BioImaging Facility (ABIF), RRID:SCR_017697.

Raw images were imported into Fiji^55,56^ for analysis. Tissue thickness quantifications in 2D, S2A, 4B, S4A, S4B and S4C were performed on images of paraffin-embedded sagittal sections using a custom Python script, available upon request. For each epidermis, the tissue was separated in bands of 30µm, the thickness of each pixel column was measured and averaged within each band to capture heterogeneity. For the average per replicate, all band thicknesses were averaged. Angles of division (3C and S3F) were measured in the Reslice view in Fiji by tracing a line through the daughter cells post-mitosis and using the basal cell layer as a reference. Hes1+ cells (1D and 1H), K10+ cells (2B), K10 reporter+ cells (S2E) and EdU+ cells (S3B) were manually quantified and expressed as a percentage of total basal or spinous cells in three 125µm x 125µm regions from separate fields of view per mouse. Total CC3+ basal cells (S3D) and total basal cells (S3E) were manually quantified in three 425µm x 425µm regions or three 125µm x 125µm regions per mouse, respectively. For longitudinal intravital imaging experiments (2G and 2H), basal cells were tracked by comparing epidermal regions imaged daily from Day 30 to Day 36 post Tamoxifen injection. Basal cells were considered to have delaminated when they could no longer be seen making any contact with the underlying basement membrane as described previously^9^. For EdU pulse-chase experiments (3G, S3I, 4F, 4G, 4H, S4E and S4F), pairs of adjacent EdU+ cells were manually scored for layer position, K10 expression, and Hes1 positivity.

## Resource table

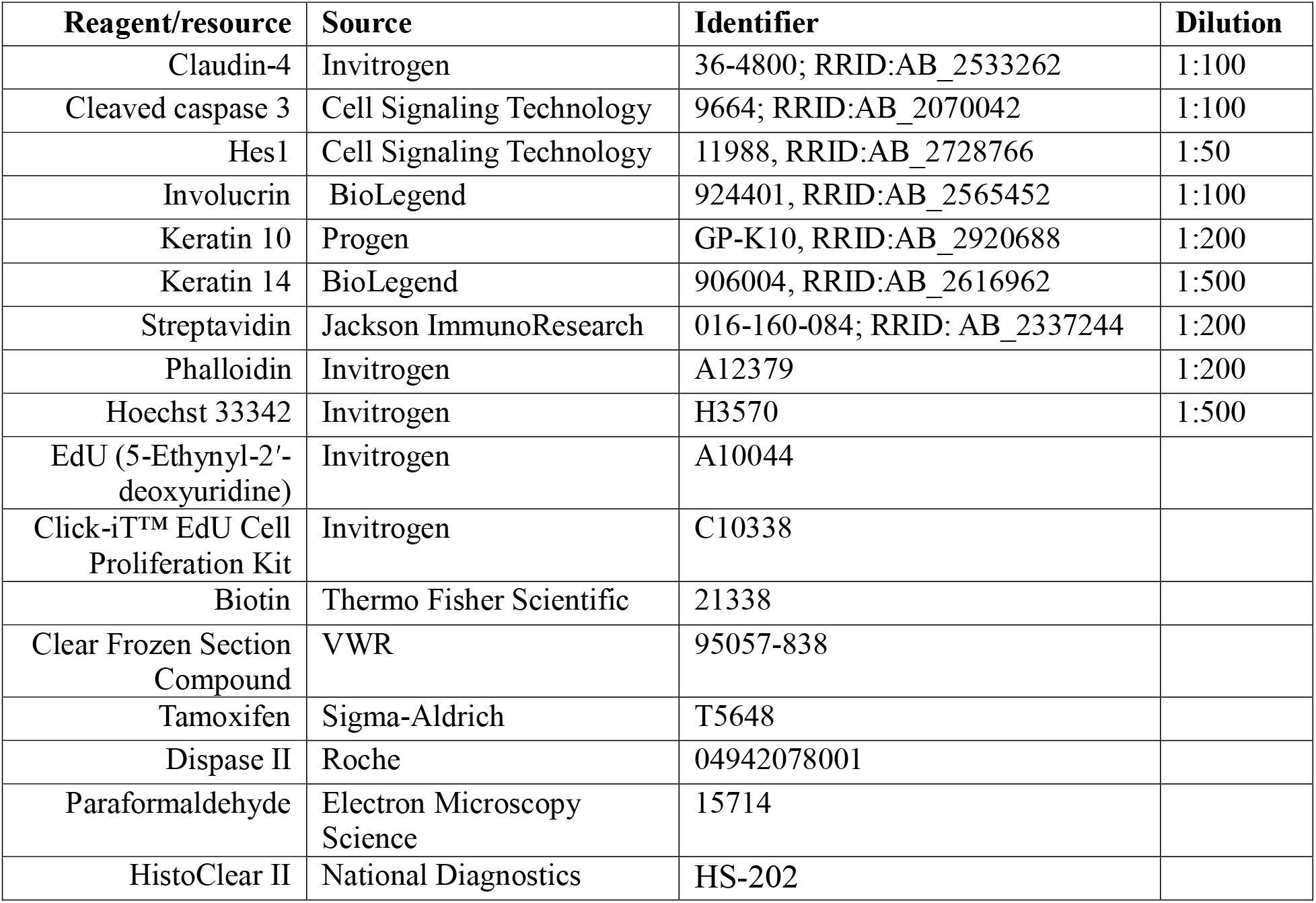

