## Supplemental files for "Jag2 patterns early differentiation in the epidermal stem cell layer"

**Affiliations:**
[1]Department of Biochemistry, McGill University, Quebec, Canada
[2]Rosalind & Morris Goodman Cancer Institute, McGill University, Canada
[3]Department of Anatomy & Cell Biology, McGill University, Canada

*These authors contributed equally

**This PDF file includes:**
Figures and Figure Legends S1 to S4

### Figure S1

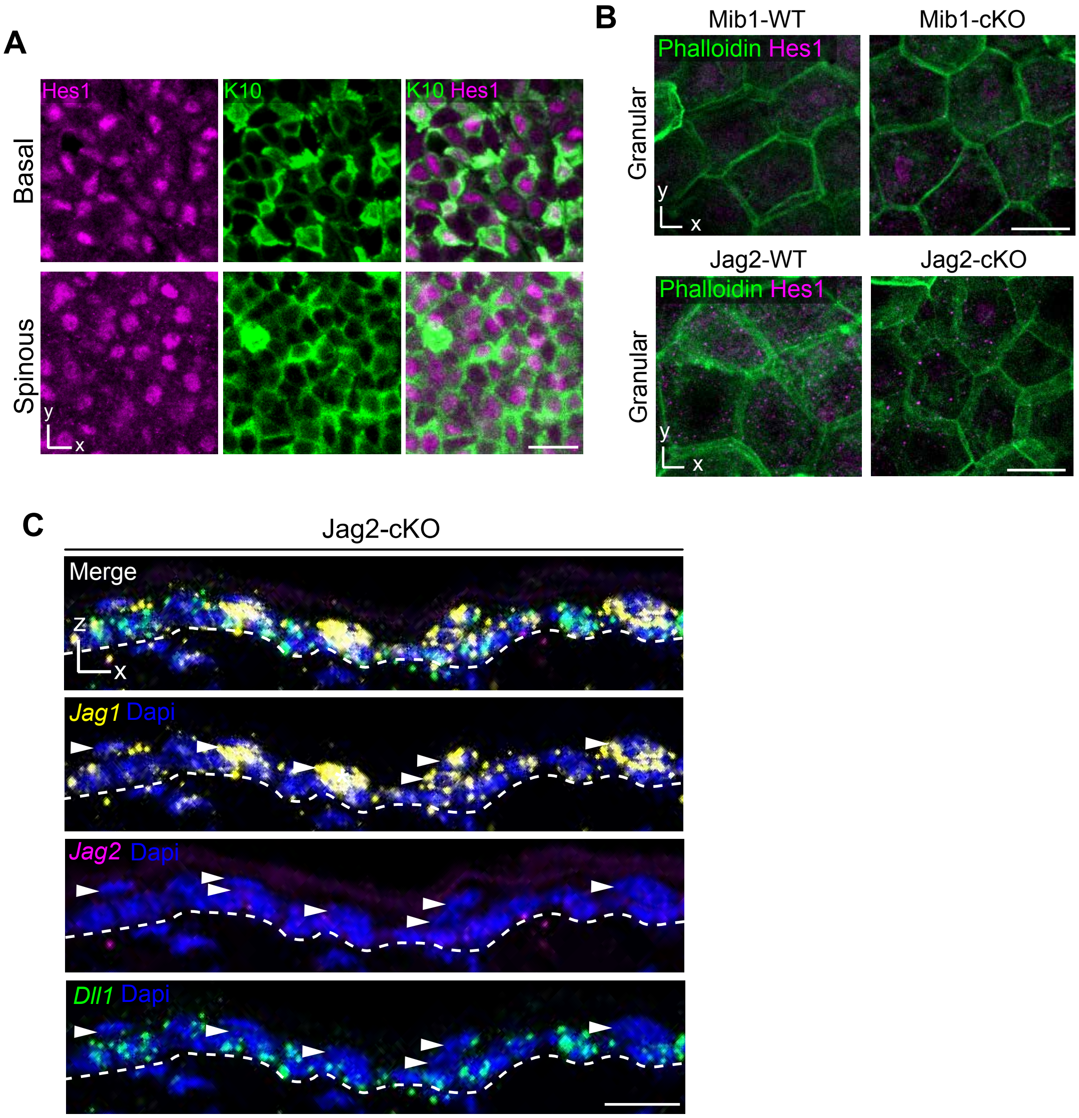


**Figure S1. Hes1 and Notch ligand expression in WT, Mib1-cKO and Jag2-cKO tissue, related to Figure 1.** (A) Whole-mount staining of Hes1 (magenta) and Keratin10 (green) in the basal and spinous layers of wildtype ear epidermis. (B) Whole-mount staining of Hes1 (magenta) in the granular layers of Mib1-WT vs Mib1-cKO (top panel) and Jag2-WT vs Jag2-cKO (bottom panel) ear epidermis. Cell boundaries are visualized with phalloidin (green). (C) Fluorescent in situ hybridization (RNAscope) of Notch ligands Jag1 (yellow), Jag2 (magenta) and Dll1 (green) with Dapi (blue) in a sagittal section of Jag2-cKO ear epidermis. White dashed line represents the basement membrane, arrowheads denote suprabasal nuclei. For images in A, B and C, Scale bar = 20μm.

### Figure S2

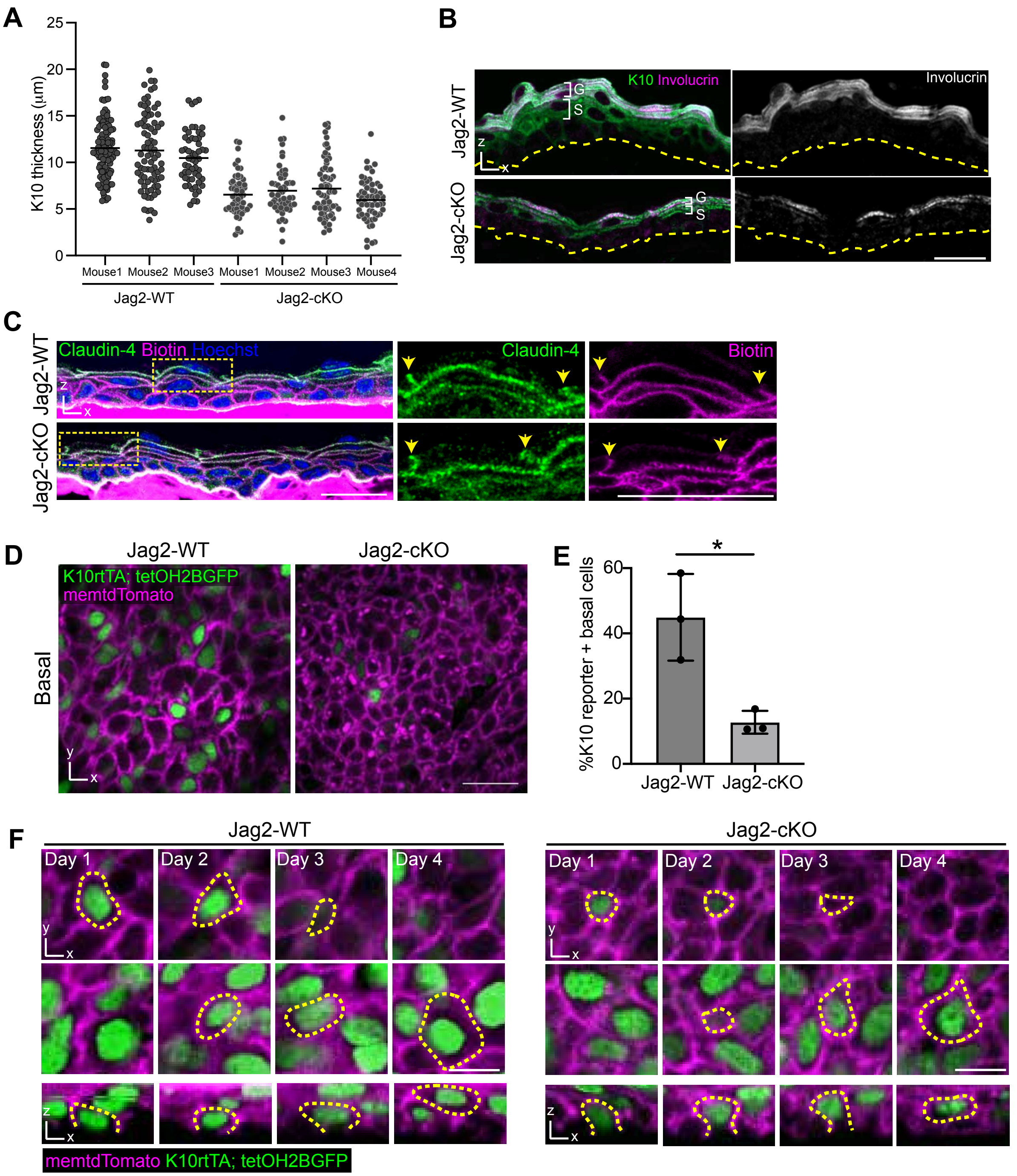


**Figure S2. Differentiation, delamination and barrier formation in Jag2-cKO tissue, related to Figure 2.** (A) Thickness of Keratin10+ layers in sagittal sections from individual Jag2-WT and Jag2-cKO mice used to generate the average values shown in Figure 2D. Individual data points represent average thickness values of 30µm wide regions. (B) Immunostaining of K10 (green) and Involucrin (magenta or white) in sagittal sections of Jag2-WT and Jag2-cKO ear epidermis. Yellow dashes represent the basement membrane. White brackets indicate boundaries of the K10 positive spinous compartment and K10/Involucrin double positive granular compartment (S and G, respectively). (C) Tight junction permeability assay performed by immunostaining of Claudin-4 (green) and intradermally-injected NHS-LC-biotin tracer (detected with streptavidin, magenta) in sagittal cryosections of Jag2-WT and Jag2-cKO ear epidermis. Dotted lines indicate inset regions (right side), and arrows highlight termination points of biotin tracer diffusion, coinciding with granular layer tight junctions. (D) Representative images of K10 reporter signal (green) in the basal layer of Jag2-WT and Jag2-cKO ear epidermis. Plasma membranes are marked with tdTomato (magenta). (E) Percentage of basal cells expressing the K10 reporter (H2BGFP+) in Jag2-WT and Jag2-cKO tissue. Bar graph represents average from n=3 mice per genotype. Student's t-test, $p \le 0.05$. (F) Longitudinal revisits of individual Jag2-WT and Jag2-cKO basal cells expressing the K10 reporter (green) across 4 days of imaging. Plasma membranes are marked with tdTomato (magenta). Yellow dotted lines indicate the same cell over time as it delaminates out of the basal layer (top row) and enters the overlying spinous layer (middle row). Bottom row shows a lateral reslice of the same cell over time. Cells are scored as having completed delamination when all contact with the basement membrane has been lost at Day 4 for both examples shown here. For images in B, C, and D, Scale bars = 20µm. For images in F, Scale bars = 10µm.

Figure S3

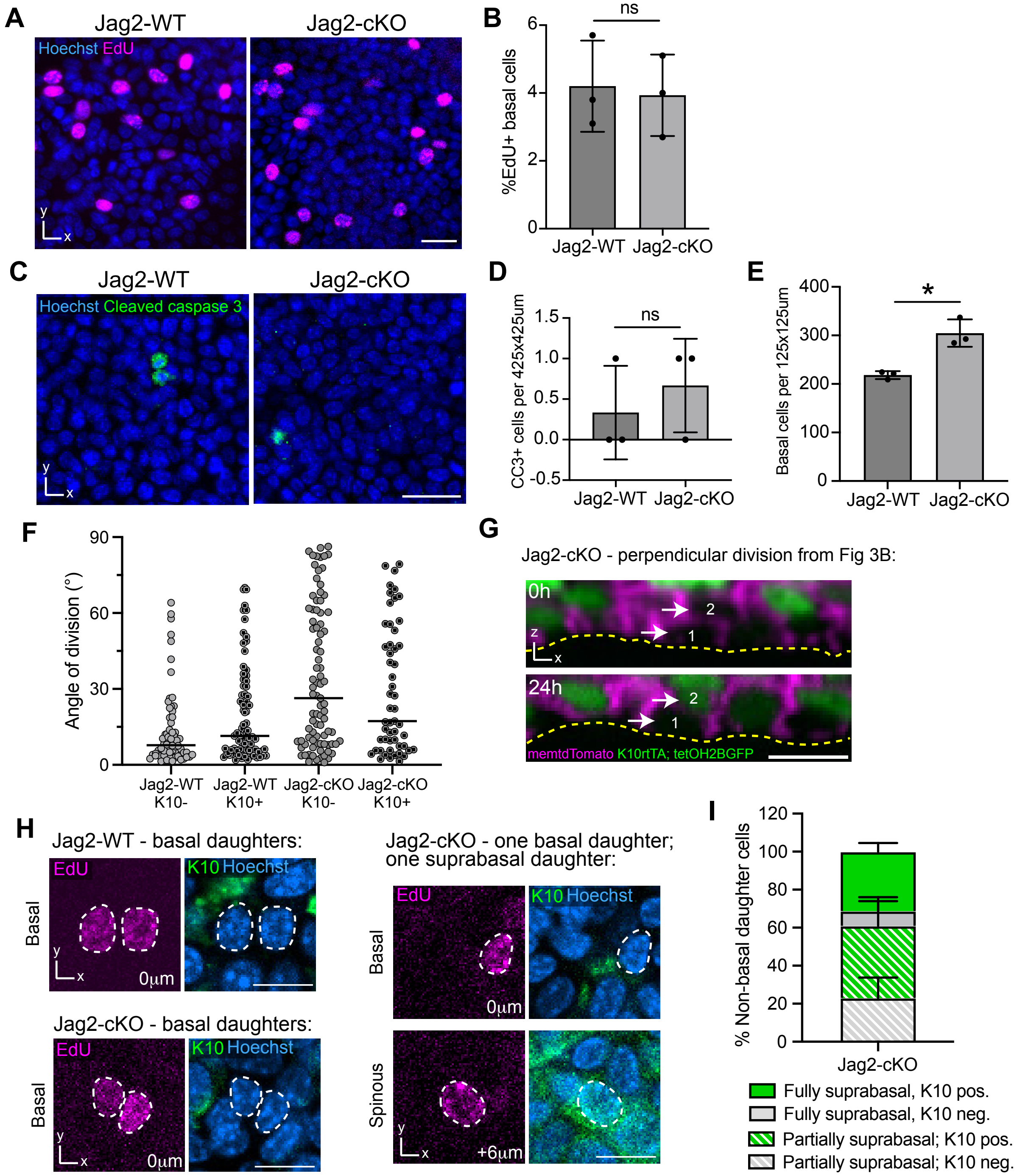


**Figure S3. Basal cell behaviors in Jag2-cKO tissue, related to Figure 3.** (A) Representative images of EdU+ cells in the basal layer 2h after EdU injection in wholemount stained Jag2-WT and Jag2-cKO tissue. Hoechst (blue) marks cell nuclei. (B) Percentage of EdU+ basal cells 2h after EdU injection in Jag2-WT and Jag2-cKO tissue. Student's t-test, $p>0.05$. (C) Whole-mount staining showing rare cleaved-caspase 3-positive (green) cells in Jag2-WT and Jag2-cKO tissue. Hoechst (blue) marks cell nuclei. (D) Quantifications of cleaved-caspase 3 (CC3)-positive cells per 425x425µm fiels of view in Jag2-WT and Jag2-cKO tissue. Student's t-test, $p>0.05$. (E) Basal cell density, expressed as the number of basal cells per 125x125µm regions in Jag2-WT and Jag2-cKO tissue. Student's t-test, $p\leq0.05$. (F) Quantification of division angle in K10 reporter positive and K10 reporter negative cells from Jag2-WT and Jag2-cKO tissue. The solid line represents the median. Graph represents n=3 mice per genotype. (G) Representative example of daughter cells generated via perpendicular division in Jag2-cKO tissue immediately following their birth (top panel) and revisited 24h later (bottom panel). Cells retain their positions in the basal and suprabasal layers over 24h (cell #1 and cell #2, respectively), and the suprabasal daughter can be seen inducing the expression of the K10 reporter. For stills from timelapse recording of this division, see Figure 3B. (H) Representative images of EdU+ daughter cells that are both basal (left images) or asymmetrically distributed in the basal and suprabasal layers (right image). Hoechst (blue) marks cell nuclei. Wholemount staining for K10 (green) shows that suprabasal daughters from Jag2-cKO tissue can initiate differentiation. (I) Percentage of EdU+ daughter cells scored as partially or fully suprabasal in Jag2-cKO tissue (see Figure 3G) that express K10. For images in A, C, G and H, scale bar = 20µm. Bar graphs in B, D, E and I represent averages from at least n=3 mice per genotype, 3 regions per mouse, error bars are mean + SD.

### Figure S4

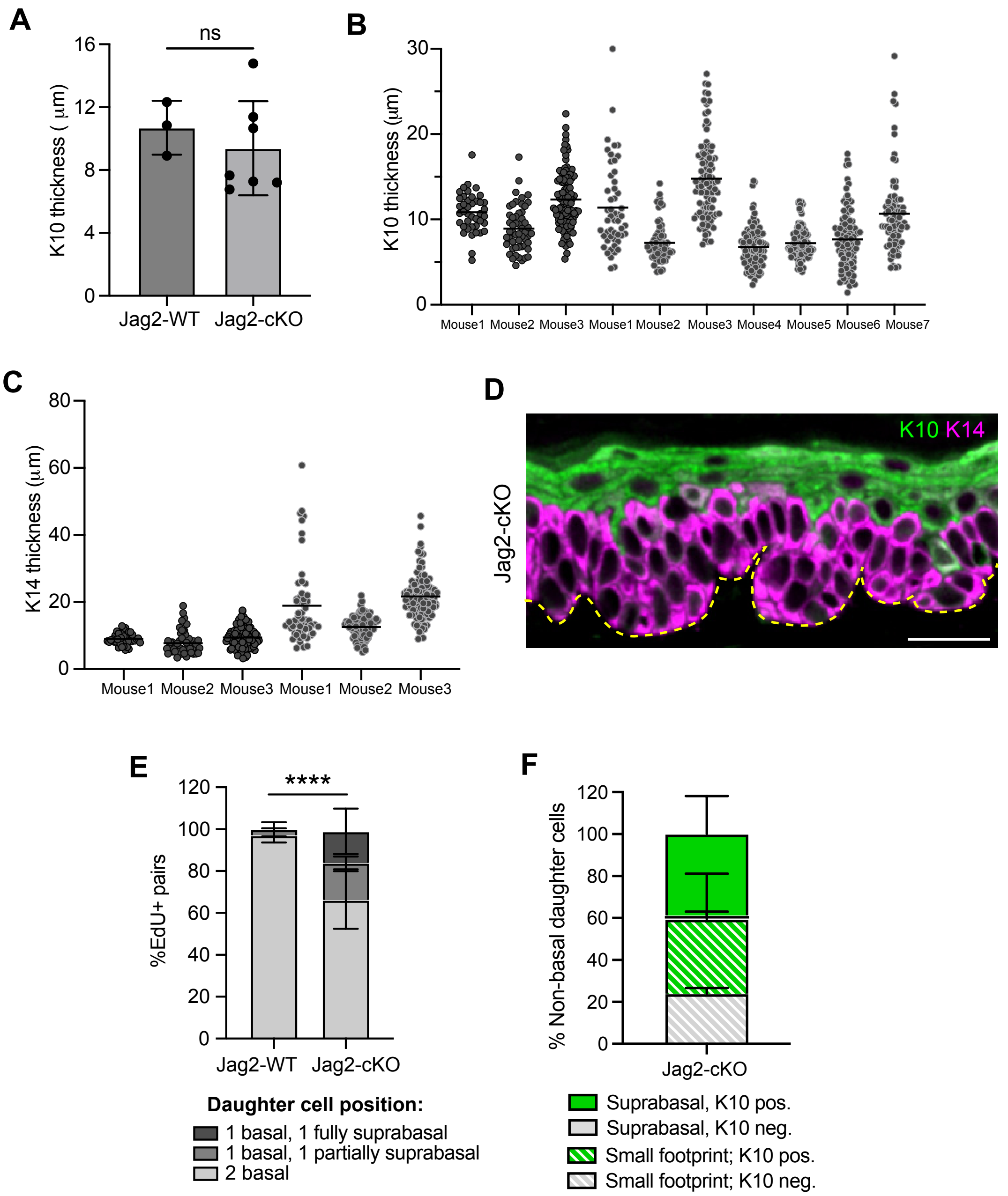


**Figure S4. Differentiation and daughter cell fate in Jag2-cKO tissue at 55 days post-deletion, related to Figure 4.** (A) Total thickness of Keratin10+ layers in sagittal sections of Jag2-WT and Jag2-cKO ear epidermis at 55 days post-deletion. Student's t-test, $p>0.05$. (B) Thickness of Keratin10+ layers in sagittal sections from individual Jag2-WT and Jag2-cKO mice used to generate the average values shown in Figure S4A. Individual data points represent average thickness values of 30µm wide regions. (C) Thickness of Keratin14+ layers in sagittal sections from individual Jag2-WT and Jag2-cKO mice used to generate the average values shown in Figure 4B. Individual data points represent average thickness values of 30µm wide regions. (D) Immunostaining of K10 (green) and K14 (magenta) in sagittal sections of Jag2-cKO tissue 6 months post-deletion, showing a severely hyperplastic K14+ basal compartment. Yellow dashes represent the basement membrane. (E) Percentage of EdU+ daughter cell pairs that are symmetrically (both basal) or asymmetrically localized (one basal and one partially/fully suprabasal) at 48h post-EdU incorporation in Jag2-WT and Jag2-cKO tissue 55 days post-deletion. Contingency tests performed per biological replicate and combined using Fisher's method; $p<0.0001$. (F) Percentage of EdU+ daughter cells scored as partially or fully suprabasal in Jag2-cKO tissue (see Figure 4E) that express K10.
